# Time spent in distinct life-history stages has sex-specific effects on reproductive fitness in wild Atlantic salmon

**DOI:** 10.1101/688572

**Authors:** Kenyon B. Mobley, Hanna Granroth-Wilding, Mikko Ellmen, Panu Orell, Jaakko Erkinaro, Craig R. Primmer

## Abstract

In species with complex life cycles, life history theory predicts that fitness is affected by conditions encountered in previous life history stages. Here, we use a four-year pedigree to investigate if time spent in two distinct life history stages has sex-specific reproductive fitness consequences in anadromous Atlantic salmon (*Salmo salar*). We determined the amount of years spent in fresh water as juveniles (freshwater age, FW), and years spent in the marine environment prior to sexual maturation (sea age, SW) on 264 spawning adults. We then estimated reproductive fitness as the number of offspring (reproductive success) and the number of mates (mating success) using genetic parentage analysis (>5000 offspring). Sea age is positively correlated with reproductive and mating success of both sexes whereby older and larger individuals gained the highest reproductive fitness benefits (females: increase of 16.5 offspring/SW and 0.86 mates/SW; males: increase of 12.4 offspring/SW and 0.43 mates/SW). Younger freshwater age was related to older sea age and thus increased reproductive fitness, but only among females (females: −9.0 offspring/FW and −0.80 mates/FW). This implies that females can obtain higher reproductive fitness by transitioning to the marine environment earlier. In contrast, male mating and reproductive success was unaffected by freshwater age and males returned to spawn earlier than females despite the fitness advantage of later sea age maturation. Our results show that the timing of transitions between juvenile and adult phases has a sex-specific consequence on female reproductive fitness, demonstrating a life-history trade-off between maturation and reproduction in wild Atlantic salmon.

## Introduction

Many organisms have complex life cycles and undergo discrete life history stages in two or more distinct habitats (Moran 1994). Transitions between these life history stages are typically accompanied by major shifts in physiology, behavior, and ecology, making them inherently risky and energetically expensive. Life history theory predicts that fitness in one life history stage may depend upon the allocation of resources in previous life history stages (Bernardo 1993; Metcalfe & Monaghan 2001; Roff 1993; Stearns 1992). To maximize fitness, tradeoffs between the duration of time spent at specific life history stages, such as the timing to switch to a new feeding habitat or when to achieve sexual maturity, are hypothesized (Roff 1993; Stearns 1992). A negative relationship between growth and the time spent in each stage may in turn affect adult fitness. For example, earlier development in one life history stage may increase the probability of surviving to reproduction (Day & Rowe 2002). However, increased survivorship may come at a cost of reduced size that could limit reproductive output through mechanisms such as decreased fecundity, increased predation, and increased mating competition (Day & Rowe 2002; Roff 2000; Stearns 1992). In sexually reproducing species, optimal strategies for growth, survival, and reproduction can also differ between the sexes (Arnqvist & Rowe 2005; Winemiller 1992). Therefore, understanding how trade-offs in the duration of time spent in specific life-history stages shape reproductive fitness between the sexes is an important next step in evolutionary biology.

Atlantic salmon (*Salmo salar*) have a complex life cycle consisting of distinct juvenile, adult, and reproductive life history stages (Jonsson & Jonsson 2011). Atlantic salmon are anadromous; sexually mature adults reproduce in fresh water, eggs hatch and juveniles stay in freshwater for a number of years prior to migrating to sea (Jonsson & Jonsson 2011). Freshwater age is the amount of time spent in the juvenile freshwater environment prior to migration to sea. The process of transitioning to seawater is known as smoltification and is associated with morphological, physiological and behavioral changes (Jonsson & Jonsson 1993; Jonsson & Jonsson 2011; McCormick *et al.* 1998). At sea, salmon spend a number of years feeding and growing at an accelerated rate prior to returning to freshwater to spawn (Fleming 1998; Fleming 1996; Jonsson & Jonsson 2011). The time spent at sea prior to returning to spawn is known as sea age and is commonly measured in sea winters (SW). Atlantic salmon exhibit wide variation in both freshwater and sea age, and this variation affects growth and the timing of reproduction (Einum *et al.* 2002; Erkinaro *et al.* 2019; Jonsson & Jonsson 1993; Jonsson & Jonsson 2011). As a result, Atlantic salmon is an excellent model system for addressing questions related to life-history evolution (Barson *et al.* 2015; Jonsson & Jonsson 2011; Stearns 1992).

A trade-off between freshwater age and sea age on Atlantic salmon reproductive fitness has been proposed by theoretical models and empirical studies (Einum *et al.* 2002; Jonsson & Jonsson 1993; Thorpe *et al.* 1998; Thorpe & Metcalfe 1998). Previous studies have shown that time spent in freshwater habitats is similar between the sexes and larger, faster growing individuals tend to spend less time in the freshwater habitat than smaller, slower growing individuals (Jonsson & Jonsson 2011; Thorpe *et al.* 1998). These individuals that spend less time in freshwater generally spend more time at sea and thus obtain sexual maturity later before returning to rivers to spawn (Erkinaro *et al.* 2019; Jonsson & Jonsson 2011). Spending more time at sea has direct reproductive fitness consequences as larger body size is related to higher fecundity (i.e., mature eggs) in females (Heinimaa & Heinimaa 2004) and higher reproductive success in both males and females (Fleming 1998; Mobley *et al.* 2019). However, spending more time at sea may come with a high cost to survivorship as fewer older individuals return to mate, presumably due to high predation at sea (McCormick *et al.* 1998; Thorpe 1994). To our knowledge, the hypothesis that a tradeoff exists between time spent in the freshwater environment and time spent at sea to maximize reproductive fitness has not yet been tested.

To date, few studies have investigated how time spent at discrete life-history stages affects reproduction in wild Atlantic salmon and previous studies focus on sea age rather than the potential for the juvenile environment to influence reproductive fitness. This is likely due, in part, to the relatively long reproductive cycle of Atlantic salmon and the low survivorship to sexual maturity in natural populations. Previous experimental studies in semi-natural settings have shown that body size is an important determinant of reproduction and is related to fecundity (numbers of eggs) in females and mate monopolization in males (Fleming *et al.* 1997; Fleming *et al.* 1996). These studies have been instrumental to our understanding of the positive relationship between body size and reproductive success in Atlantic salmon but were conducted in controlled settings using populations with limited life history variation (e.g. wild salmon males all 1 SW, females 1-2 SW, (Fleming *et al.* 1997) and did directly measure reproductive success via genetic parentage reconstruction. Garant *et al.* (2001, 2003), also found evidence for a relationship between body size, reproductive success and mating success in Atlantic salmon in 1SW and 2SW males and 2SW females, but did not characterize freshwater age, rendering comparisons between concerning life history stages incomplete. Building on more recent developments in genetic techniques, sex-specific life-history trade-offs can measure the reproductive success of spawning adults by reconstructing pedigrees over multiple years (e.g., Christie *et al.* 2018; Mobley *et al.* 2019).

In this study, we use data from Mobley *et al.* (2019) to dig deeper into how sex-specific effects of timing of two major life-history stages, freshwater age and sea age, may affect reproductive fitness. The optimal time spent in the freshwater and marine environment may differ between male and female Atlantic salmon in order to maximize reproductive fitness. The dataset used consists of parentage analysis on 264 adults with life history information and >5000 juveniles collected for over four-cohort years from a population of wild Atlantic salmon from Northern Finland. We first tested for a sex-specific relationship between freshwater age and sea age to see if time in freshwater affects the time spent in seawater differently between the sexes. Second, we investigated the relationship between adult body size and freshwater age and sea age to determine whether time spent at these life history stages affected the overall size at reproduction. Third, we tested whether males and females differed in the relationship between reproductive success and mating success, which can inform us on whether sexual selection is acting stronger on males or females. Finally, we looked for sex-specific differences in the relationship between effect of freshwater age and sea age on reproductive and mating success. The nature of these relationships were first tested using a complete dataset with all adults including those that did not have offspring in our sample (all adults). We also tested these relationships using reduced datasets that only included breeding adults (breeding adults) and only first-time spawning adults excluding repeat-spawning individuals (first-time spawners).

## Methods

Anadromous adults were sampled in September-October 2011-2014 at the lower Utsjoki spawning grounds at the mouth of the Utsjoki tributary of the Teno River in northern Finland (69°54’28.37’’N, 27°2’47.52’’E, see Mobley *et al.* (2019) for further details on sampling location). Fishing permission for research purposes was granted by the Lapland Centre for Economic Development, Transport, and the Environment (permit numbers 1579/5713-2007, 2370/5713-2012, and 1471/5713-2017). Adults were primarily captured by gill nets at night to minimize handling stress. A few males were captured by rod and reel angling. Adults were sexed, weighed, and total length was recorded. Condition was calculated as the residual from a linear model of weight predicted by length for each sex and spawning cohort (Mobley *et al.* 2019; Patterson 1992). Scales were collected for age analysis and a small piece of anal fin was collected for genetic analysis prior to release near the site of capture. Juveniles were sampled by electrofishing shallow areas in the region of the spawning grounds 10 to 11 months later, which is two to three months after they are expected to have emerged from the nests in the stream bed gravel (Mobley *et al.* 2019). Genetic samples were collected from all juveniles by collecting a small piece of adipose and/or anal fins, after which they were immediately returned to the river (Mobley *et al.* 2019). Four parent-offspring cohorts were sampled in this manner between 2011 and 2015.

### Age determination

Freshwater age, defined as the number of years spent in freshwater prior to migrating to sea, and sea age, defined as the number of years an individual overwintered at sea before returning to spawn, was determined for adults captured on the spawning ground using scale growth readings as outlined in Aykanat *et al.* (2015). Freshwater age could not be determined on 25 individuals (3 females, 22 males) using scale data. Sea age could not be accomplished for 16 adults > 1SW (1 female, 15 males) using scale data. Extrapolating sea age based on calculated distributions of weight of known sea age individuals could be accomplished for all adults missing sea age (see Mobley *et al.* 2019, Supplementary Materials, Table S4). However, freshwater age was not extrapolated based on weight due to the poor relationship between weight and freshwater age (see Results). Therefore, these individuals were excluded from statistical analyses. Repeat spawners that were spawning for a second time were also determined using scale data. Thirteen individuals (6 females, 7 males) were identified as repeat spawners by scale aging analysis. The mean sea age at maturity of repeat spawning females was 3.2 ± 0.4 SE (range 2-4 SW) and all repeat spawning males had spent one year at sea before the first spawning migration and another year at sea before returning to spawning for the second time (i.e., all male repeat spawners were 2 SW).

### Parentage analysis

Molecular parentage analysis was conducted according to Mobley *et al.* (2019). Briefly, all adults and juveniles were genotyped using 13 microsatellite loci previously used for parentage analyses in this species (Aykanat *et al.* 2014). Pedigrees were constructed for each parent-offspring cohort separately using the package MasterBayes V2.55 (Hadfield *et al.* 2006) in the R programming environment (R Core Team 2018) with conditions as outlined in Mobley *et al.* (2019).

### Reproductive fitness estimates

Reproductive success was quantified as the number of offspring assigned to an adult, following parentage assignment of all offspring. Mating success was estimated as the number of unique mates per individual identified within our sample by parentage analysis (Mobley *et al.* 2019).

### Statistical analyses

We tested for a sex difference in the relationship between freshwater age and sea age and their interaction using a linear regression model. Sex differences in the relationship between weight, body size (length), and condition, and their interaction with sex were tested in linear models for freshwater age and sea age separately. We also tested for a sex difference in the relationship between reproductive success and mating success using a linear regression model.

We tested for a sex difference in the relationship between reproductive success and mating success using a generalized linear mixed model (GLMM) approach fitting separate models for freshwater age and sea age. All GLMM models of reproductive and mating success included an offset of the number of offspring sampled in the relevant year, log-transformed for consistency with the models’ link functions to account for between-year variation in sampling effort. We applied zero-inflated models using the function zeroinfl() from the package *pscl* (Jackman 2017; Zeileis *et al.* 2008) because estimates of mating success and reproductive success contained a high proportion of individuals without any offspring, and hence mates, assigned. These models were mixture models that accounted for zero values in both of its two parts: a binomial model for the frequency of zeros and, conditional on this, a count model using a Poisson distribution. Effects of freshwater and sea age were tested only in the count part of the model, but both count and binomial parts included an offset to account for differences in sampling effort between years. For reproductive success, an effect of sex was also included in the binomial part because a much greater proportion of males did not have any sampled offspring compared to females. For mating success, sex was not a significant predictor in the binomial part and hence was not included in the final models.

All response variables for models were first tested with a full initial model consisting of an interaction between sex and the relevant explanatory variable and their main effects for all adults (all adult datset). Non-significant (p > 0.05) explanatory variables were removed step-wise from the model to obtain a minimal model in which all predictors had a significant effect. In all models, the four sampling years were pooled to maximize sample sizes as patterns in reproductive success have been shown to be consistent across years (Mobley *et al.* 2019).

Models of weight, length, condition, reproductive success and mating success were also tested on two additional reduced datasets. The “breeding adults”, consisting only of those individuals that were assigned offspring in our sample (non-zero number of offspring and mates). Because repeat spawning may influence reproductive fitness, the breeding adults dataset was further reduced to only include “first time spawners” thereby excluding the 13 repeat spawners. For the GLMM on reproductive success and mating success, reduced datasets were modelled using a negative binomial general linear model (GLM) in the R package MASS for reproductive success, and using a quasipoisson GLM in the R package stats for the number of mates. Distributions were chosen based on the behaviour of model residuals. All statistical models were performed in R (R Core Team 2018) and all means are reported ± one standard error of the mean.

## Results

A total of 230 adult males, 34 adult females and 5223 juvenile offspring (<1 year old) were collected over the four cohort years. At the time of spawning, females were larger, heavier and had older sea age than males, on average (Table 1). However, both sexes had similar condition estimates at spawning (Table 1). Means and sample sizes for weight, body size, condition, freshwater age, sea age, reproductive success and mating success for all adults, breeding adults (adults with offspring assigned from our sample), and first-time spawners (excluding repeat-spawners) pooled across cohort years are reported in Table 1 and summarized by freshwater age and sea age in Table S1.

**Table 1.** Summary of length, age and reproductive fitness estimates for all adults pooled across cohort years, only breeding adults, and first-time spawners (breeding adults without respawners). The number of adults (*n*) and mean weight (kg), length (cm), condition, freshwater age (FW age, years), sea age (sea winters, SW), reprodutive success (No. offspring) and mating success (No. mates) for each sex are given.

|  | *n* | Weight (kg) | Length (cm) | Condition | FW age* (years) | Sea age (SW) | No. offspring | No. mates |
| --- | --- | --- | --- | --- | --- | --- | --- | --- |
| **All adults** |  |  |  |  |  |  |  |  |
| *Females* | 34 | 7.61 ± 0.63 | 89.2 ± 2.3 | 0.00 ± 0.19 | 3.58 ± 0.12 | 2.41 ± 0.13 | 26.5 ± 6.4 | 2.47 ± 0.36 |
| *Males* | 230 | 3.65 ± 0.21 | 70.0 ± 0.9 | 0.00 ± 0.05 | 3.65 ± 0.04 | 1.35 ± 0.05 | 6.1 ± 1.0 | 0.71 ± 0.06 |
| **Breeding adults** |  |  |  |  |  |  |  |  |
| *Females* | 28 | 8.23 ± 0.66 | 91.6 ± 2.1 | 0.01 ± 0.18 | 3.54 ± 0.13 | 2.54 ± 0.14 | 32.2 ± 7.4 | 3.00 ± 0.37 |
| *Males* | 115 | 4.54 ± 0.39 | 74.1 ± 1.6 | -0.04 ± 0.09 | 3.63 ± 0.06 | 1.53 ± 0.08 | 12.3 ± 1.8 | 1.43 ± 0.07 |
| **First-time spawners** |  |  |  |  |  |  |  |  |
| *Females* | 22 | 7.43 ± 0.59 | 89.2 ± 1.9 | -0.17 ± 0.14 | 3.57 ± 0.15 | 2.41 ± 0.13 | 27.0 ± 6.3 | 3.05 ± 0.43 |
| *Males* | 111 | 4.49 ± 0.40 | 73.8 ± 1.7 | -0.03 ± 0.09 | 3.64 ± 0.06 | 1.51 ± 0.08 | 12.2 ± 1.9 | 1.44 ± 0.07 |

\*Freshwater age *n* (males, females): all adults = 31, 211; breeding adults = 26, 102; first time spawners = 21, 89).

Females spent less time, on average, in the juvenile freshwater environment than males prior to migrating to sea (freshwater age: −0.45 ± 0.62, *t*_1,235_ = −4.105, p < 0.0001, Table 1, Fig. 1). Mean sea age, on the other hand, was higher in females than males (sex: −1.16 ± 0.16, *t*_1,235_ = −7.374, p <0.0001; Table 1, Fig. 1). We found a sex-specific effect between freshwater age and sea age: older freshwater age females returned to spawn at younger sea ages, whereas freshwater age had no effect on sea age in males (sex* freshwater age: 0.41 ± 0.18, *t*_1,235_ = 2.343, p = 0.0200; Fig. 1). However, only the sex difference in sea age was significant when restricting analyses to breeding adults and first-time spawner datasets (Table S2).

**Fig. 1.**
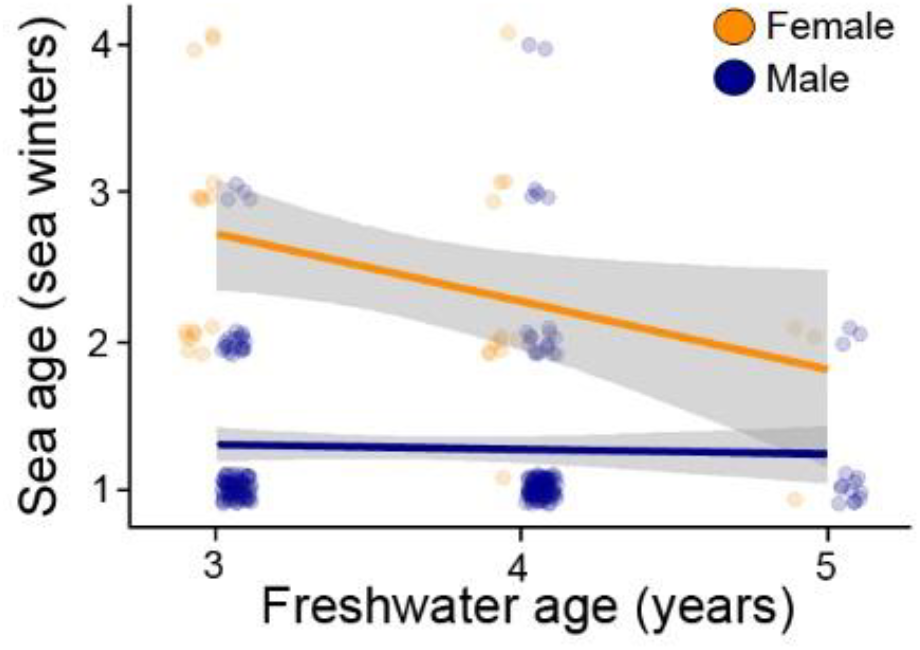
Sex difference in the relationship between freshwater age and sea age in the “all adults dataset”. Colored lines represent linear regression for each sex and gray areas represent 95% confidence intervals (CI). Small circles show individual data points. For clarity, individual points are jittered on the x and y axis.

Mirroring the negative sex-specific relationship between freshwater age and sea age, females that spent more time in freshwater suffered a significant decrease in weight at sexual maturity (sex: −10.63 ± 3.04, *t*_1,238_ = −3.497, p = <0.0006; freshwater age: −1.721 ± 0.77, *t*_1,238_ = −2.242, p = <0.0259; sex*freshwater age: 1.70 ± 0.83, *t*_1,238_ = 2.037, p = 0.0427; Fig. 2a). However, only the sex difference in weight was significant while restricting analyses to the breeding adults and first-time spawner datasets (Table S3). Adults that spent more time at sea were heavier and no sex difference was discerned (sex: −0.09 ± 1.35, *t*_1,261_ = −0.066, p = 0.9470; sea age: 17.98 ± 0.64, *t*_1,261_ = 27.988, p = <0.0001; Fig. 2b). When restricting the analyses to the breeding adults and first-time spawners, significant effects of both sex and sea age were apparent (Table S3).

**Fig. 2.**
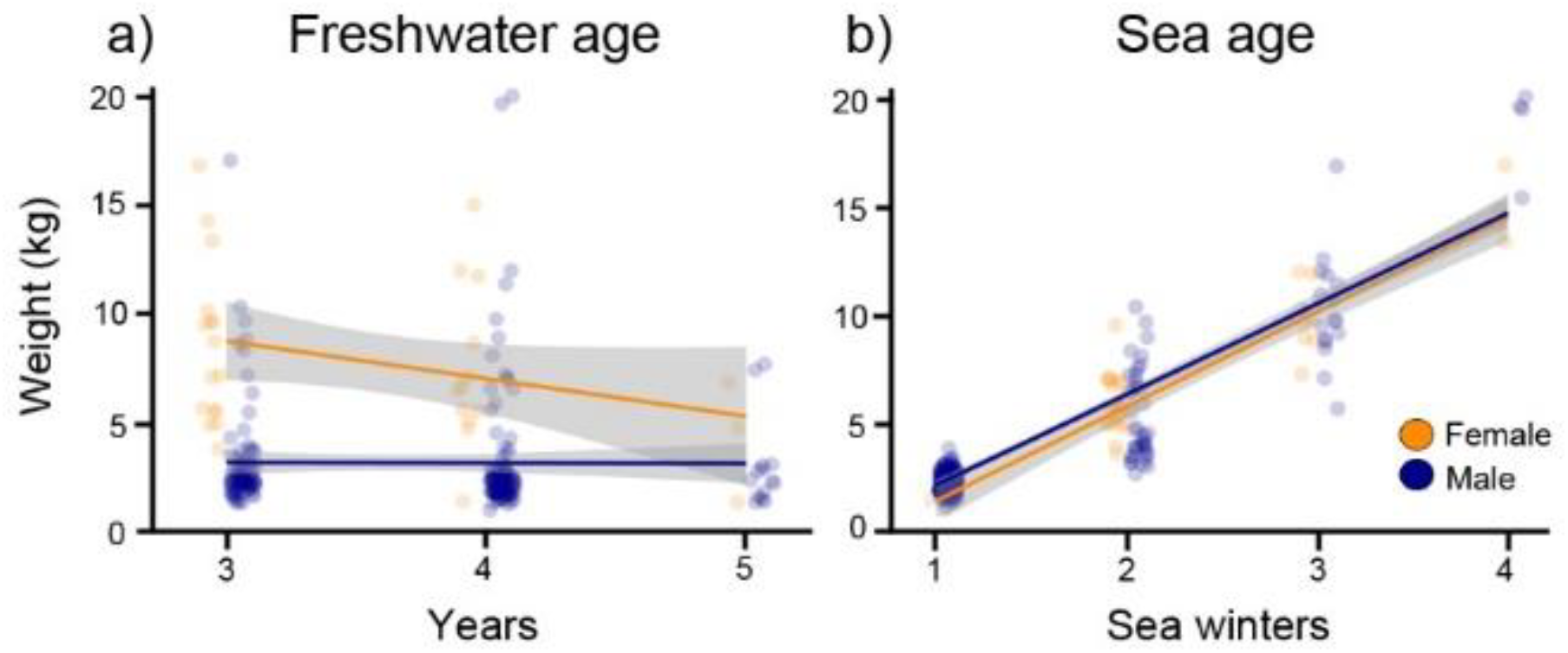
The influence of a) freshwater and b) sea age on weight at reproductive maturity for males and females in the “all adults dataset”. Colored lines represent linear regression for each sex and gray areas represent 95% confidence intervals (CI). Small circles show individual data points. For clarity, individual points are jittered on the x and y axis.

Freshwater age did not affect body size in either females or males (sex: −21.46 ± 2.30, *t*_1,239_ = −9.320, p = <0.0001; freshwater age: −2.28 ± 1.27, *t*_1,239_ = 1.800, p = 0.0731). Sea age, as expected, was positively and significantly related to body size, and this relationship was similar between the sexes (sex: −0.09 ± 1.89, *t*_1,245_ = −0.066, p = 0.947; sea age: 17.98 ± 0.64, *t*_1,245_ = 27.988, p < 0.0001; sex*sea age, 2.77 ± 1.49, *t*_1,244_ = 1.855, p = 0.0649). Relationships between length and freshwater age and sea age were similar when analyzing only breeding adults or first time spawners with the exceptions that sex differences in body size and a significant interaction between sex and sea age was observable (Table S4).

Freshwater age did not affect overall condition in either females or males (sex: −0.13 ± 0.15, *t*_1,239_ = −0.862, p = 0.3890; freshwater age: 0.06 ± 0.82, *t*_1,239_ = 0.669, p = 0.5040). Similarly, sea age did not affect condition at spawning (sex: 0.12 ± 0.18, *t*_1,261_ = 0.666, p = 0.5060; sea age: 0.11 ± 0.08, *t*_1,261_ = 1.476, p = 0.1410). However, condition was significantly higher in higher sea-age individuals in the restricted breeding adult and first-time spawner datasets (Table S5).

Bayesian parentage analysis of parents and juveniles using 13 microsatellite loci assigned 1987 of the offspring (38%) to at least one sampled adult with confidence. Based on estimates of the number of offspring and mates assigned to adults from parentage analysis, females, on average, had higher reproductive and mating success than males. Females had a mean 26.5 ± 6.5 offspring (range 0-177) and 2.47 ± 0.36 mates (range 0-8, Table 1). Females gained an average of 5.10 ± 1.56 offspring/kg (*t*_1,32_ = 10.68, P < 0.0026) and 0.22 ± 0.09 mates/kg (*t*_1,32_ = 5.73 3, P = 0.0227, Fig. S1). Males had a mean of 6.1 ± 1.0 offspring (range 0-145) and 0.71 ± 0.05 mates (range 0-5, Table 1). Males gained an average of 2.87 ± 0.25 offspring/kg (*t*_1,228_ = 134.33, P < 0.0001) and 0.10 ± 0.02 mates/kg (*t*_1,228_ = 32.80, P < 0.0001; Fig. S1).

The nature of the relationship between reproductive success and mating success was similar between males and females (mates: 11.25 ± 0.83, *t*_1,261_ = 13.557, p < 0.0001; sex: −0.62 ± 3.11, *t*_1,261_ = −0.199, p = 0.842; sex*mates, −3.065 ± 1.658, *t*_1,260_ = −1.849, p = 0.0656; Fig. 3). These results were also similar when analyzing only breeding adults or first time spawners (Table S6).

**Figure 3.**
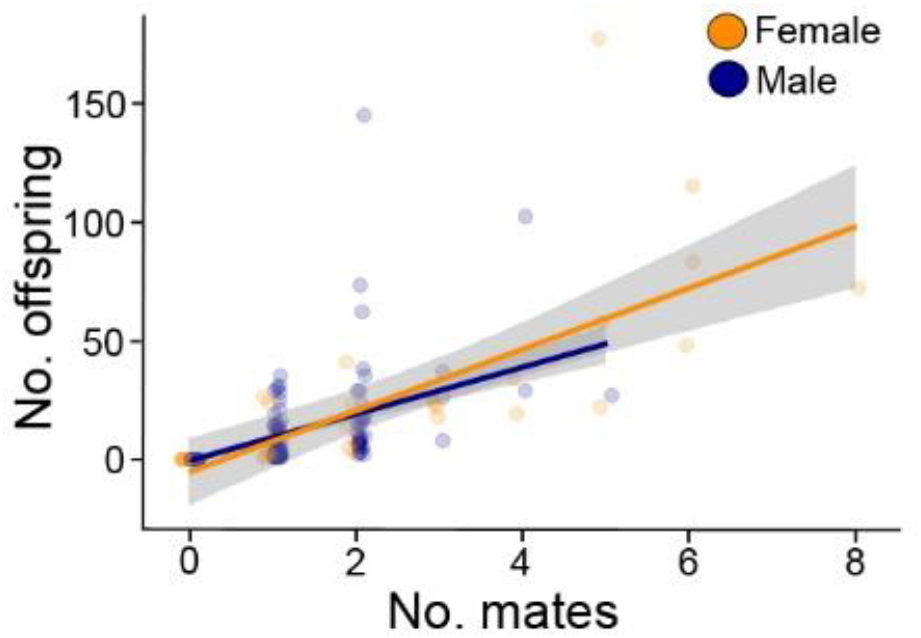
The relationship of reproductive success No. offspring) with mating success (No. mates) for male and females in the “all adults dataset”. Colored lines represent linear regression for each sex and gray areas represent 95% confidence intervals (CI). Small circles show individual data points. For clarity, individual points are jittered on the x and y axis.

There was a negative relationship between freshwater age and reproductive and mating success in females (−9.01 ± 10.60 offspring/FW, *t*_1,29_ = −0.85, p = 0.4024; −0.80 ± 0.57 mates/FW, *t*_1,29_ = −1.40, p = 0.1720). In contrast, no relationship between reproductive and mating success and freshwater age in males was found (0.52 ± 1.48 offspring/FW, *t*_1,209_ = 0.35, p = 0.7286, 0.05 ± 0.09 mates/FW, *t*_1,209_ = 0.57, p = 0.5676). Both reproductive success and mating success showed positive relationships with sea age in females, although only reproductive success was significant (16.53 ± 7.99 offspring/SW, *t*_1,32_ = 2.07, p = 0.0466; 0.86 ± 0.45 mates/SW, *t*_1,32_ = 1.90, p = 0.0671). Both reproductive and mating success was positively correlated to sea age in males (12.45 ± 1.19 offspring/SW, *t*_1,228_ = −5.91, p < 0.0001; 0.44 ± 0.08 mates/SW, *t*_1,228_ = 5.54, p <0.0001).

Results of GLMM minimal models demonstrated that more time spent in the juvenile freshwater habitat was associated with reduced reproductive success, but only in females (Table 2; Fig. 4a). However, adults that spent more time at sea had greater reproductive success, with a steeper relationship in females, and females had greater reproductive success overall (Table 2, Fig. 4b). However, when restricting the dataset to breeding adults and first time spawners, no effect of freshwater age on reproductive success was significant whereas the effect of sea age and sex remained significant (Table S7).

**Table 2.** Results of GLMM minimal models showing the effect of sex differences on freshwater (FW) age and sea age and on reproductive success and mating success. Effect sizes for age are shown per year, and for sex, for males compared to females, with the response transformed according to the relevant link function. Freshwater age, sea age, and sex are count data whereas zero inflation terms are binomial.

|  | Effect size | SE | z | Pr(>\|z\|) |
| --- | --- | --- | --- | --- |
| **Reproductive success** |  |  |  |  |
| *Freshwater age* |  |  |  |  |
| Intercept | -2.22 | 0.20 | -10.85 | <0.0001 |
| FW age | -0.40 | 0.06 | -6.86 | <0.0001 |
| Sex | -3.61 | 0.28 | -13.09 | <0.0001 |
| FW age*Sex | 0.64 | 0.08 | 8.33 | <0.0001 |
| Zero inflation (Intercept) | -8.82 | 0.49 | -17.96 | <0.0001 |
| Zero inflation (Sex) | 1.64 | 0.51 | 3.21 | 0.0010 |
| *Sea age* |  |  |  |  |
| Intercept | -5.12 | 0.12 | -44.12 | <0.0001 |
| Sea age | 0.57 | 0.04 | 14.07 | <0.0001 |
| Sex | -1.09 | 0.13 | -8.18 | <0.0001 |
| Sea age*Sex | 0.23 | 0.05 | 4.78 | <0.0001 |
| Zero inflation (Intercept) | -8.74 | 0.45 | -19.21 | <0.0001 |
| Zero inflation (Sex) | 1.46 | 0.47 | 3.09 | 0.0020 |
| **Mating success** |  |  |  |  |
| *Freshwater age* |  |  |  |  |
| Intercept | -4.39 | 0.71 | -6.18 | <0.0001 |
| FW age | -0.49 | 0.20 | -2.37 | 0.0180 |
| Sex | -3.78 | 0.90 | -4.22 | <0.0001 |
| FW age*Sex | 0.66 | 0.25 | 2.61 | 0.0090 |
| Zero inflation (Sex) | -9.15 | 0.48 | -19.21 | <0.0001 |
| *Sea age* |  |  |  |  |
| Intercept | -7.21 | 0.24 | -29.96 | <0.0001 |
| Sea age | 0.42 | 0.08 | 5.41 | <0.0001 |
| Sex | -0.93 | 0.16 | -5.75 | <0.0001 |
| Zero inflation (Sex) | -9.41 | 0.53 | -17.14 | <0.0001 |

**Figure 4.**
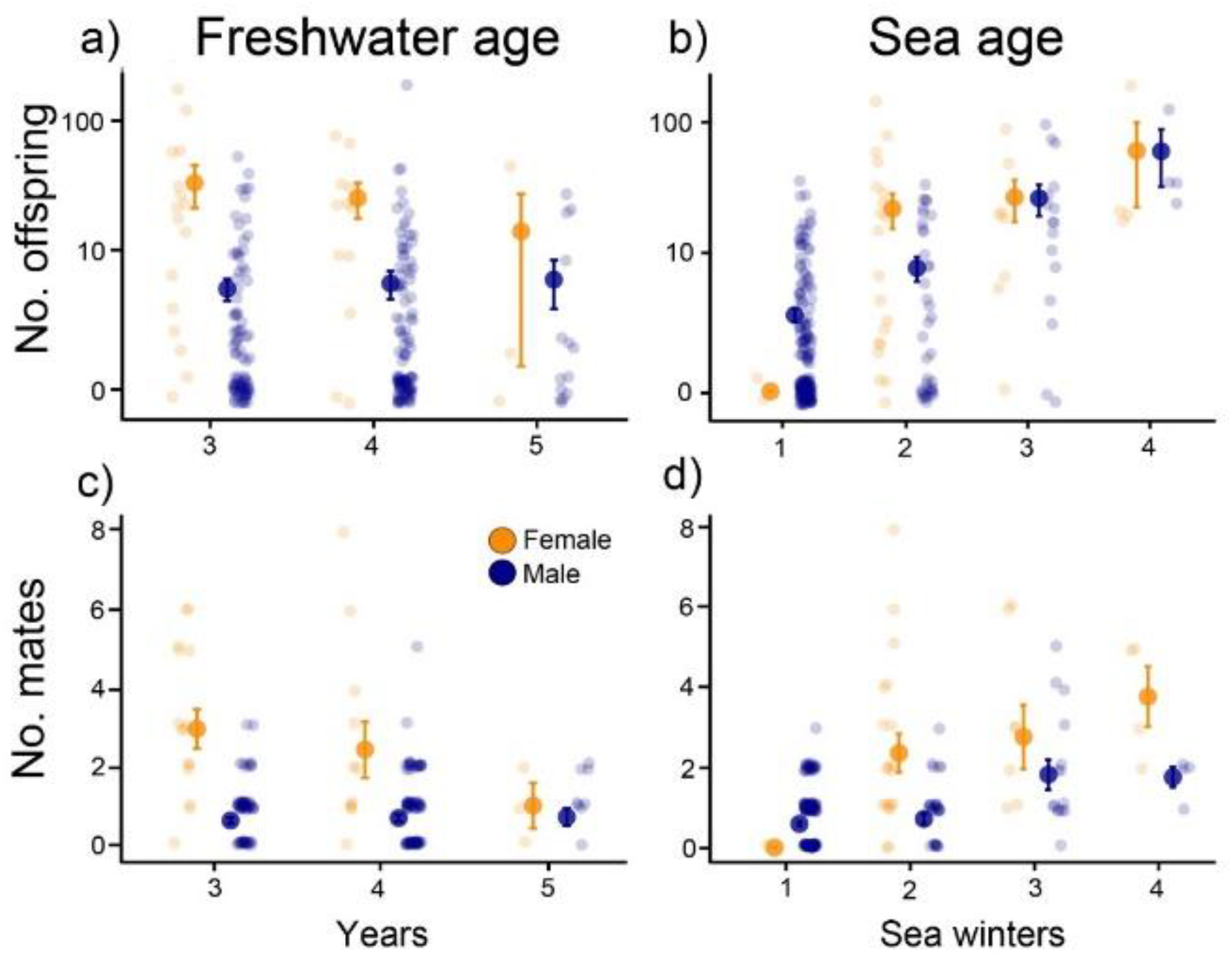
The relationship of reproductive success (No. offspring) with a) freshwater age and b) sea age, and mating success (No. mates) with c) freshwater age and d) sea age in male and female Atlantic salmon in the “all adults dataset”. Large circles with error bars represent the mean ± SE, small circles show individual data points. For clarity, individual points are jittered on the x and y axis and the y axis for reproductive success are log_10_ transformed.

Patterns in mating success mirrored those in reproductive success. Females that spent longer in the juvenile freshwater habitat showed reduced mating success, while males’ mating success was not affected by freshwater age (Table 2, Fig. 4c). A general increase in the number of mates with sea age in both males and females was observed (Table 2, Fig. 4d). All of these effects remained significant within in the restricted datasets among breeding adults and first time spawners (Table S4).

## Discussion

In this study, we investigated sex-specific trade-offs in reproductive fitness and the time spent during two life-history stages of anadromous Atlantic salmon. A sex-specific trade-off between time spent in the freshwater stage and reproductive fitness was apparent among females. Females that remained longer in freshwater spent less time at sea before returning to spawn, were smaller, suffering a slight, but significant, reduction in both reproductive and mating success. In contrast, males spent less time at sea than females but showed no indication that freshwater age influenced reproductive fitness. The time spent at sea had a substantial positive influence on weight, body size, and reproductive fitness of both sexes and condition among breeding adults. Moreover, any negative effect of longer time spent in freshwater on reproductive fitness in females is masked by the strong positive relationship between sea age, body weight/size, and reproductive fitness.

In our study, females were larger and spent more years at sea than males, as is the case for most populations of Atlantic salmon (Barson *et al.* 2015). Females also had more mates and produced more offspring than males. We found a significant positive relationship between mating and reproductive success in both male and female Atlantic salmon indicating that having more mating partners results in more offspring in both sexes similar to reports in North American Atlantic salmon (Garant *et al.* 2001). The relationship between reproductive success and mating success does not differ between the sexes suggesting that the strength of sexual selection is similar between males and females (Anthes *et al.* 2017; Arnold & Duvall 1994; Janicke *et al.* 2016; Jones 2009). This result is surprising, as it is generally thought that sexual selection is stronger among males in Atlantic salmon (Fleming 1998; Fleming 1996; Fleming & Einum 2011). We expected the relationship to be greater in males as there was a significant 7:1 male-bias in the sex ratio of adults caught on the spawning grounds that was consistent over the four cohort years (Mobley *et al.* 2019). Based on mating system theory, a male-biased sex ration should drive higher levels of mate competition among males for available females (Emlen & Oring 1977; Mobley 2014; Shuster & Wade 2003). Potentially, high levels of sneaking by younger anadromous males and mature male parr (pre-smolting individuals that have not yet transitioned to the marine environment) can decrease sexual selection among males (e.g., Jones *et al.* 2001). However, we should be careful with this interpretation as no information on reproductive success of these parr in our study population is currently available but, occurrence of mature male parr in the region is relatively low (Heinimaa & Erkinaro 2004). Further research is warranted to uncover the extent to which sexual selection operates on males and females in this and indeed other species of salmonids (Auld *et al.* 2019).

The strong positive relationship between sea age and weight, body size, and reproductive fitness estimates in both sexes is likely related to genetic and environmental factors controlling sea age at maturation. Sea age is partially under genetic control of the *vgll3* locus in this population (Ayllon *et al.* 2015; Barson *et al.* 2015; Czorlich *et al.* 2018) explaining nearly 40% of the variation in sea age. This same genomic region may also influence the potential for repeat-spawning (iteroparity) in this species (Aykanat *et al.* 2019). Environmental factors may also affect sea age including salinity, photoperiod, and temperature (Fjelldal *et al.* 2011; Melo *et al.* 2014). Compared to our understanding of the genetic and environmental underpinnings of sea age, the conditions responsible for the timing of smoltification in Atlantic salmon are less well understood. The decision to leave the freshwater juvenile environment likely depends upon the balance between growth and survival at sea (McCormick *et al.* 1998; Thorpe 1994). Earlier smolting individuals spend more time at sea where they are potentially exposed to higher predation (McCormick *et al.* 1998). Previous studies do not appear to show clear patterns concerning a fitness trade-off between freshwater age and sea age. For example, a negative relationship between smolt size, pre-smolt growth and post-smolt growth was reported earlier in female Atlantic salmon from Norway (Einum *et al.* 2002), yet no relationship between mean growth and sea age at maturity was found in Spanish Atlantic salmon (Nicieza & Braña 1993). Other species of salmonids, steelhead trout (*O. mykiss*) and coho salmon (*O. kisutch*), show a weak positive association between pre- and post-smolt growth indicating no trade-off between freshwater age and sea age in these species, at least under artificial hatchery conditions (Johnsson *et al.* 1997). Environmental conditions may also affect smolt timing as smoltification is also associated with warmer water temperatures (Duston & Saunders 1997). Currently, it is unknown whether genes associated with *vgll3* affect freshwater age, and future studies should investigate genetic and environmental factors underpinning sex differences in smolt timing in an effort to understand their relative contributions to reproductive fitness.

In our study, We were able to assign 37.5% offspring to at least one sampled adult confidently (Mobley *et al.* 2019, Supplementary Materials). Because it is often difficult to recover all breeding individuals and offspring in large, open, natural populations, missing parentage data can potentially bias estimates of mating and reproductive success (Mobley 2014; Mobley & Jones 2013). For example, adults could have produced offspring but were not recovered in our sample would have their overall contributions to reproduction underestimated. However, the results of our zero-inflated models for male and female mating and reproductive success that accounted for individuals that did not have offspring in our sample were generally supported by analyses that excluded individuals that did not produce offspring in our sample (breeding adults) and excluding multi-year spawners (first time spawners) (Table S7). With the exception that freshwater age did not influence mating and reproductive success when individuals that did not have offspring in our sample were excluded, all other comparisons showed similar results using breeding adults and first time spawners. These results demonstrate that our analyses are generally robust to the exclusion of these individuals.

This study complements an earlier study concerning local adaptation in the Teno river salmon (Mobley *et al.* 2019). In the earlier study, a small proportion (12.5% of the adults) were identified as dispersers (i.e., strays) from genetically distinct populations within the Teno River system. Reproductive success among local females and males was 9.6. and 2.9 times higher than among dispersers, respectively. Our choice to include these dispersers in this study reflects the reality that these adults were present on the spawning grounds and a substantial portion of dispersers, particularly females, produced offspring, albeit with lower success. Moreover, restricting the analyses to only breeding adults (Table S7) did not change the overall results relating to mating and reproductive success indicating that inclusion of potentially transient individuals did not appreciably affect our overall conclusions. However, the inclusion of such dispersers in this study may influence our interpretations in two ways. First, inclusion of dispersers reduces the mean reproductive fitness estimates as dispersers have demonstratively lower mating and reproductive success than locals. Second, dispersers originating from different tributaries may be locally adapted to different environmental conditions that affect freshwater age. Further, dispersers may have been still migrating upstream to spawn although we believe that this is an unlikely scenario for reasons outlined in Mobley *et al.* (2019).

Despite the reduced reproductive fitness of individuals that return after only one year at sea, it is possible that these individuals will gain further reproductive fitness by repeat-spawning. Although the majority of individuals spawn after only one SW, Atlantic salmon are iterparous, and both males and females can return to spawn over multiple years (Hutchings & Morris 1985; Jonsson & Jonsson 2011; Niemelä *et al.* 2006). A recent study has shown in a pacific salmon, *O. mykiss*, repeat-spawning individuals may obtain 2.5 times the lifetime reproductive success than single spawners (Christie *et al.* 2018). Furthermore, it has recently been shown that the potential for repeat spawning in Atlantic salmon is associated with the *vgll3* locus and is tied to the decision to return earlier from sea to spawn for the first time (Aykanat *et al.* 2019). However, only a small number of repeat spawners were captured on the spawning grounds and our results were robust to the exclusion of these individuals (Table S7). Thus, we lack sufficient data to address this topic presently and hope that it can be analyzed in the future with the addition of more cohort years.

## Conclusion

A fundamental goal of evolutionary biology is to understand how life-history trade-offs affect individual fitness. This study contributes to this goal by investigating reproductive fitness of the timing of transitions at two critical life-history stages and demonstrating that there is a sex-specific life-history trade-off between maturation and reproduction in a wild population of anadromous salmon. Indirect costs may also play a role in life-history stages, as early smolting individuals may be at greater risk of mortality via predation at sea yet may also have a higher chance of multiple reproductive seasons. Future research should investigate sex-specific growth rates and the timing of smoltification, as well as mortality at sea to uncover the potential hidden costs associated with maturation at specific life-history stages.

## Acknowledgments

We thank Katja Salminen, Meri Lindqvist, Jenni Kuismin, Jani Aaltonen, Susanna Ukonaho, and Jan Laine for laboratory assistance, Jorma Kuusela, Jari Haantie and Matti Kylmäaho for scale aging analyses, and Olavi Guttorm, Topi Pöyhönen, Timo Kanniainen, Arto Koskinen, Jorma Ollila, Mari Lajunen, Tuomo Karjalainen (deceased), Mikko Kytökorpi, Seda Karslioglu, Anna Ellmen and Hans Pieski for field assistance. We thank Tutku Aykanat and Paul Debes for fruitful discussions.

## Author contributions

K.B.M. & C.R.P. conceived the study. M.E., P.O and J.E. co-ordinated and/or participated in sample collection. C.R.P co-ordinated the molecular data generation. K.B.M. and H.G.-W. analyzed the data. K.B.M. drafted the manuscript, with input from all other authors.

## Data and materials availability

All data will be uploaded to the DRYAD digital repository upon publication of the article.

## Supplementary materials

Figure S1, Tables S1-7.

