## Supplementary Materials for "Time spent in distinct life-history stages has sex-specific effects on reproductive fitness in wild Atlantic salmon"

**This file includes:**

Fig S1

Tables S1 to S7

**Figure S1.** Reproductive success (# offspring, RS) and mating success (# mates, MS) as a function of body weight. Numbers indicate age, either freshwater or sea age, of females (orange) and males (blue). Freshwater age could not be assigned to 25 individuals

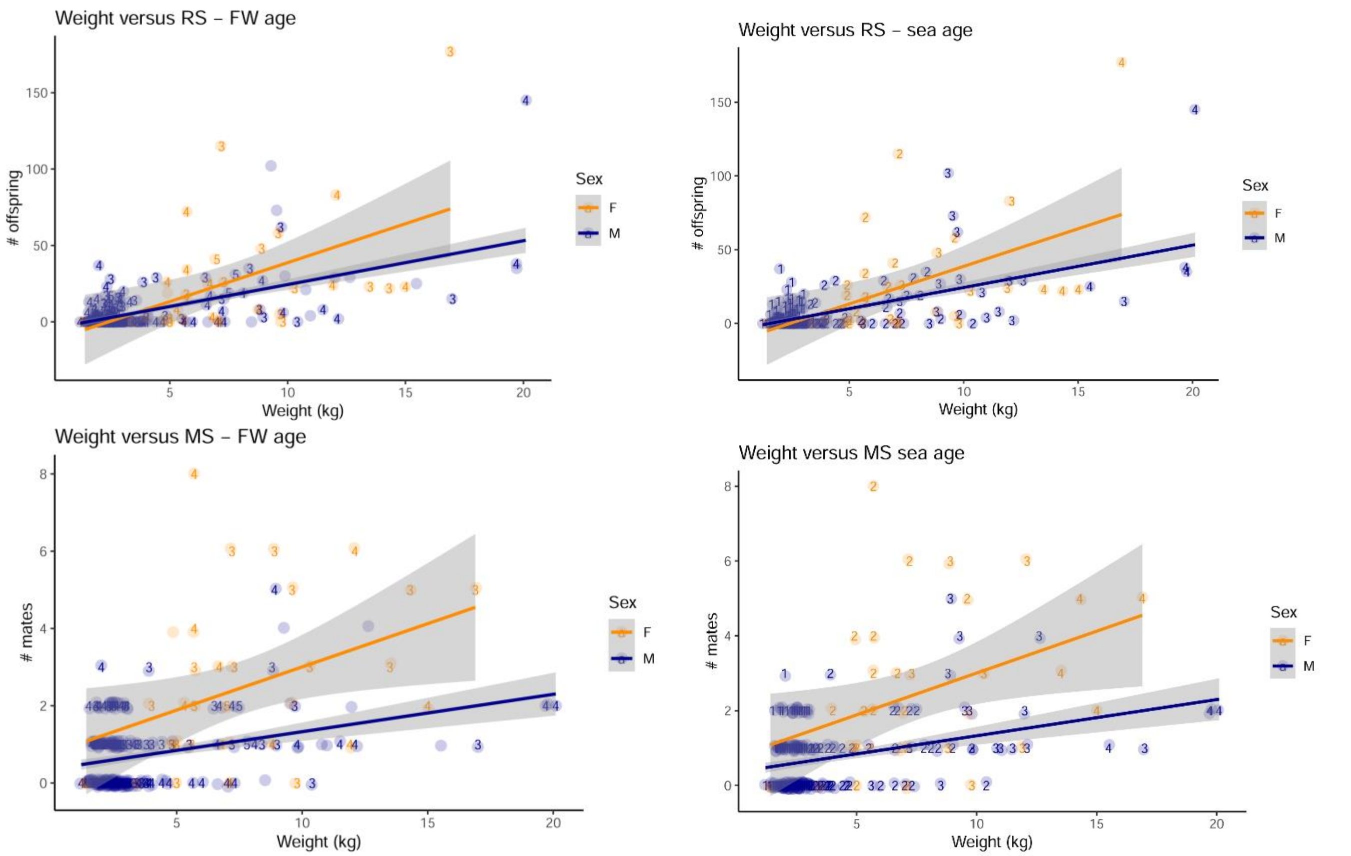

**Table S1**. Summary of data for freshwater age (FW age) and sea age (in seawinters, SW). Number of individuals (n), means (± SE) for weight, body length, condition, reproductive success (# offspring), and mating success (#mates) are listed (all adults dataset).

| | Freshwater age (FW) | | | | Sea age (SW) | | | |
|---|---|---|---|---|---|---|---|---|
| | 3 | 4 | 5 | | 1 | 2 | 3 | 4 |
| Females | | | | | | | | |
| n | 16 | 12 | 3 | | 2 | 20 | 8 | 4 |
| Weight (kg) | 8.66 ± 0.93 | 7.63 ± 1.08 | 4.4 ± 1.63 | | 1.45 ± 0.05 | 5.87 ± 0.31 | 9.85 ± 0.57 | 14.92 ± 0.72 |
| Length (cm) | 93.6 ± 3.1 | 88.5 ± 3.9 | 74.3 ± 9.9 | | 56.5 ± 0.5 | 84.3 ± 1.1 | 98.3 ± 1.3 | 112.0 ± 1.22 |
| Condition | -0.07 ± 0.26 | 0.16 ± 0.30 | 0.65 ± 0.94 | | 2.06 ± 0.33 | -0.38 ± 0.15 | -0.18 ± 0.30 | 1.25 ± 0.89 |
| # offspring | 33.4 ± 12.1 | 25.5 ± 7.8 | 14.0 ± 13.5 | | -- | 22.1 ± 6.7 | 27.0 ± 9.6 | 61.3 ± 38.6 |
| # mates | 2.94 ± 0.50 | 2.41 ± 0.71 | 1.0 ± 0.58 | | -- | 2.35 ± 0.48 | 2.75 ± 0.80 | 3.75 ± 0.75 |
| Males | | | | | | | | |
| n | 88 | 109 | 14 | | 172 | 38 | 16 | 4 |
| Weight (kg) | 3.28 ± 0.25 | 3.32 ± 0.29 | 3.08 ± 0.54 | | 2.31 ± 0.04 | 5.35 ± 0.34 | 10.30 ± 0.65 | 18.8 ± 1.09 |
| Length (cm) | 69.15 ± 1.20 | 68.1 ± 1.3 | 67.5 ± 3.2 | | 63.4 ± 0.3 | 81.4 ± 1.7 | 101.7 ± 2.0 | 120.8 ± 2.2 |
| Condition | -0.06 ± 0.07 | -0.01 ± 0.07 | -0.11 ± 0.14 | | 0.09 ± 0.03 | -0.69 ± 0.12 | -0.17 ± 0.30 | 3.48 ± 1.05 |
| # offspring | 4.6 ± 1.0 | 5.2 ± 1.4 | 5.6 ± 2.6 | | 2.7 ± 0.4 | 7.3 ± 1.7 | 26.6 ± 7.3 | 60.8 ± 28.2 |
| # mates | 0.63 ± 0.08 | 0.69 ± 0.09 | 0.71 ± 0.22 | | 0.59 ± 0.06 | 0.71 ± 0.12 | 2.81 ± 0.37 | 1.75 ± 0.25 |

**Table S2.** Linear models describing sex differences in sea age (SW) as a function of freshwater age (FW) for adults, breeding adults and first-time spawners datasets. Bold p values indicate significance (α = 0.05).

| | Effect size | SE | *t* | p |
|---|---|---|---|---|
| **All adults** | | | | |
| SW ~ Sex * FW | | | | |
| Intercept | 4.071 | 0.598 | 6.813 | **<0.0001** |
| Sex | -2.669 | 0.651 | -4.105 | **<0.0001** |
| FW | -0.452 | 0.164 | -2.756 | **0.0063** |
| Sex:FW | 0.417 | 0.178 | 2.343 | **0.0200** |
| **Breeding adults** | | | | |
| Sea age ~ Sex + FW | | | | |
| Intercept | 3.034 | 0.392 | 7.749 | **<0.0001** |
| Sex | -1.162 | 0.158 | -7.374 | **<0.0001** |
| FW | -0.129 | 0.103 | -1.250 | 0.2130 |
| **First-time spawners** | | | | |
| SW ~ Sex + FW | | | | |
| Intercept | 2.814 | 0.396 | 7.105 | **<0.0001** |
| Sex | -1.043 | 0.167 | -6.259 | **<0.0001** |
| FW | -0.108 | 0.103 | -1.053 | 0.2940 |

**Table S3.** Linear models for sex differences in body weight as a function of freshwater age (FW) and sea age (SW) for adults, breeding adults and first-time spawners datasets. Bold p values indicate significance (α = 0.05).

| | Effect size | SE | *t* | p |
|---|---|---|---|---|
| **All adults** | | | | |
| Weight ~ Sex * FW | | | | |
| Intercept | 14.006 | 2.795 | 5.012 | **<0.0001** |
| Sex | -10.633 | 3.041 | -3.497 | **<0.0006** |
| FW | -1.721 | 0.767 | -2.242 | **0.0259** |
| Sex:FW | 1.697 | 0.833 | 2.037 | **0.0427** |
| Weight ~ Sex + SW | | | | |
| Intercept | -2.611 | 0.372 | -7.019 | **<0.0001** |
| Sex | 0.513 | 0.281 | 1.827 | 0.0689 |
| SW | 4.238 | 0.120 | 35.312 | **<0.0001** |
| **Breeding adults** | | | | |
| Weight ~ Sex + FW | | | | |
| Intercept | 9.401 | 1.947 | 4.829 | **<0.0001** |
| Sex | -4.439 | 0.784 | -5.664 | **<0.0001** |
| FW | -0.283 | 0.514 | -0.551 | 0.5830 |
| Weight ~ Sex + SW | | | | |
| Intercept | -3.290 | 0.477 | -6.891 | **<0.0001** |
| Sex | 0.874 | 0.349 | 2.506 | **0.0133** |
| SW | 4.543 | 0.152 | 29.852 | **<0.0001** |
| **First-time spawners** | | | | |
| Weight ~ Sex + FW | | | | |
| Intercept | 7.914 | 1.970 | 4.016 | **<0.0001** |
| Sex | -3.681 | 0.829 | -4.438 | **<0.0001** |
| FW | 0.103 | 0.510 | -0.201 | 0.8409 |
| Weight ~ Sex + SW | | | | |
| Intercept | -3.581 | 0.470 | -7.624 | **<0.0001** |
| Sex | 1.159 | 0.354 | 3.274 | **0.0014** |
| SW | 4.570 | 0.150 | 30.393 | **<0.0001** |

**Table S4.** Linear models for sex differences in body length as a function of freshwater age (FW) and sea age (SW) for adults, breeding adults and first-time spawners datasets. Bold p values indicate significance (α = 0.05).

| | Effect size | SE | *t* | p |
|---|---|---|---|---|
| **All adults** | | | | |
| Length ~ Sex + FW | | | | |
| Intercept | 96.877 | 5.229 | 18.528 | **<0.0001** |
| Sex | -21.108 | 2.403 | -8.782 | **<0.0001** |
| FW | -1.993 | 1.319 | -1.319 | 0.132 |
| Length ~ Sex + SW | | | | |
| Intercept | 44.828 | 1.594 | 28.11 | **<0.0001** |
| Sex | 0.229 | 1.203 | 0.19 | 0.849 |
| SW | 18.401 | 0.514 | 35.78 | **<0.0001** |
| **Breeding adults** | | | | |
| Length ~ Sex + FW | | | | |
| Intercept | 101.677 | 7.937 | 12.811 | **<0.0001** |
| Sex | -20.378 | 3.195 | -6.378 | **<0.0001** |
| FW | -2.691 | 2.094 | -1.295 | 0.2010 |
| Length ~ Sex * SW | | | | |
| Intercept | 56.592 | 3.995 | 14.165 | **<0.0001** |
| Sex | -12.294 | 4.154 | -2.960 | **0.0036** |
| SW | 13.795 | 1.514 | 9.112 | **<0.0001** |
| Sex:SW | 5.676 | 1.648 | 3.443 | **<0.0001** |
| **First-time spawners** | | | | |
| Length ~ Sex + FW | | | | |
| Intercept | 96.806 | 8.183 | 11.830 | **<0.0001** |
| Sex | -18.286 | 3.444 | -5.310 | **<0.0001** |
| FW | -2.039 | 2.118 | -0.963 | 0.3380 |
| Length ~ Sex * SW | | | | |
| Intercept | 56.472 | 5.022 | 11.245 | **<0.0001** |
| Sex | -12.176 | 5.134 | -2.372 | **0.0192** |
| SW | 13.578 | 2.027 | 6.697 | **<0.0001** |

| | | | | |
|---|---|---|---|---|
| Sex:SW | 5.929 | 2.118 | 2.799 | **0.0059** |

**Table S5.** Linear models for sex differences in condition as a function of freshwater age (FW) and sea age (SW) for adults, breeding adults and first-time spawners datasets. Bold p values indicate significance (α = 0.05).

| | Effect size | SE | *t* | p |
|---|---|---|---|---|
| **All adults** | | | | |
| Condition ~ Sex + FW | | | | |
| Intercept | -0.108 | 0.326 | -0.331 | 0.7410 |
| Sex | -0.129 | 0.150 | -0.862 | 0.3890 |
| FW | 0.055 | 0.822 | 0.669 | 0.5040 |
| Condition ~ Sex + SW | | | | |
| Intercept | -0.268 | 0.233 | -1.148 | 0.2520 |
| Sex | 0.117 | 0.176 | 0.666 | 0.5060 |
| SW | 0.111 | 0.075 | 1.476 | 0.1410 |
| **Breeding adults** | | | | |
| Condition ~ Sex + FW | | | | |
| Intercept | -0.637 | 0.473 | -1.346 | 0.1810 |
| Sex | -0.153 | 0.190 | -0.801 | 0.4240 |
| FW | 0.186 | 0.125 | 1.489 | 0.1390 |
| Condition ~ Sex + SW | | | | |
| Intercept | -0.912 | 0.302 | -3.023 | **0.0030** |
| Sex | 0.313 | 0.220 | 1.418 | 0.1583 |
| SW | 0.363 | 0.096 | 3.772 | **0.0002** |
| **First-time spawners** | | | | |
| Condition ~ Sex + FW | | | | |
| Intercept | -8.030 | 0.463 | -1.734 | 0.0855 |
| Sex | 0.031 | 0.195 | 0.159 | 0.8741 |
| FW | 0.186 | 0.120 | 1.548 | 0.1244 |
| Length ~ Sex + SW | | | | |
| Intercept | -0.986 | 0.305 | -3.229 | **0.0016** |
| Sex | 0.447 | 0.230 | 1.941 | 0.0544 |
| SW | 0.339 | 0.098 | 3.466 | **0.0007** |

**Table S6.** Linear models for sex differences in reproductive success (No. offspring) as a function of mating success (No. mates) for adult, breeding adult and first-time spawner datasets. Bold p values indicate significance (α = 0.05).

| | Effect size | SE | *t* | p |
|---|---|---|---|---|
| **All adults** | | | | |
| No. Offspring ~ Sex * No. mates | | | | |
| Intercept | -5.362 | 3.945 | -1.359 | 0.1753 |
| Sex | 4.483 | 4.143 | 1.082 | 0.2803 |
| No. mates | 12.908 | 1.219 | 10.587 | **<0.0001** |
| Sex * No. mates | -3.065 | 1.658 | -1.849 | 0.0656 |
| No. offspring ~ Sex + No. mates | | | | |
| Intercept | -1.265 | 3.280 | -0.386 | 0.7000 |
| Sex | -0.618 | 3.106 | -0.199 | 0.8420 |
| No. mates | 11.250 | 0.830 | 13.557 | **<0.0001** |
| **Breeding adults** | | | | |
| No. offspring ~ Sex * No. mates | | | | |
| Intercept | -9.295 | 7.053 | -1.318 | 0.190 |
| Sex | 3.997 | 8.217 | 0.486 | 0.627 |
| No. mates | 13.837 | 1.978 | 6.996 | **<0.0001** |
| Sex * No. mates | -1.511 | 3.303 | -0.457 | 0.648 |
| No. offspring ~ Sex + No. mates | | | | |
| Intercept | -7.670 | 6.075 | -1.263 | 0.209 |
| Sex | 0.989 | 4.914 | 0.201 | 0.841 |
| No. mates | 13.295 | 1.580 | 8.416 | **<0.0001** |
| **First-time spawners** | | | | |
| No. offspring ~ Sex * No. mates | | | | |
| Intercept | -8.6890 | 6.8770 | -1.263 | 0.209 |
| Sex | 2.8297 | 7.8329 | 0.361 | 0.718 |
| No. mates | 11.7188 | 1.8909 | 6.198 | **<0.0001** |
| Sex * No. mates | 0.8211 | 2.9991 | 0.274 | 0.785 |
| No. offspring ~ Sex + No. mates | | | | |
| Intercept | -9.683 | 5.820 | -1.664 | 0.0986 |
| Sex | 4.537 | 4.724 | 0.960 | 0.336 |
| No. mates | 12.045 | 1.462 | 8.236 | **<0.0001** |

**Table S7.** Results for zero-inflated mixture models (GLMM) showing the effect of sex differences in freshwater age (FW) and sea age (SW) on reproductive success and mating success for breeding adults and first-time spawners. Bold p values indicate significance (α = 0.05). All age*sex interactions were included in the initial model but removed if not significant.

| | Effect size | SE | *z* | p |
|---|---|---|---|---|
| **REPRODUCTIVE SUCCESS** | | | | |
| **FW** | | | | |
| *Breeding adults* | | | | |
| No. offspring ~ Sex + FW | | | | |
| Intercept | -3.82 | 0.60 | -6.32 | **<0.0001** |
| Sex | -1.37 | 0.24 | -5.72 | **<0.0001** |
| FW | 0.06 | 0.16 | 0.40 | 0.688 |
| *First time spawners* | | | | |
| No. offspring ~ Sex + FW | | | | |
| Intercept | -4.36 | 0.63 | -6.96 | **<0.0001** |
| Sex | -1.23 | 0.26 | -4.76 | **<0.0001** |
| FW age | 0.16 | 0.16 | 1.01 | 0.311 |
| **SW** | | | | |
| *Breeding adults* | | | | |
| No. offspring ~ Sex + SW | | | | |
| Intercept | -5.43 | 0.31 | -17.25 | **<0.0001** |
| SW | 0.69 | 0.10 | 6.86 | **<0.0001** |
| Sex | -0.62 | 0.23 | -2.72 | **0.0006** |
| *First-time spawners* | | | | |
| No. offspring ~ Sex + SW | | | | |
| Intercept | -5.46 | 0.32 | -16.99 | **<0.0001** |
| SW | 0.70 | 0.10 | 6.77 | **<0.0001** |
| Sex | -0.62 | 0.24 | -2.59 | **0.0097** |
| **MATING SUCCESS** | | | | |
| **FW** | | | | |
| *Breeding adults* | | | | |
| No. mates ~ Sex * FW | | | | |
| Intercept | -4.64 | 0.53 | -8.682 | **<0.0001** |
| FW | -0.40 | 0.15 | -2.609 | 0.0102 |

| | | | | |
|---|---|---|---|---|
| Sex | -3.12 | 0.67 | -4.693 | **<0.0001** |
| FW*Sex | 0.61 | 0.19 | 3.298 | **0.0013** |
| *First-time spawners* | | | | |
| No. mates ~ Sex * FW | | | | |
| Intercept | -4.86 | 0.59 | -8.122 | **<0.0001** |
| FW age | -0.34 | 0.17 | -2.026 | **0.0451** |
| Sex | -2.90 | 0.73 | -3.996 | **0.0001** |
| FW*Sex | 0.56 | 0.20 | 2.776 | **0.0064** |
| **SW** | | | | |
| *Breeding adults* | | | | |
| No. mates ~ Sex + SW | | | | |
| Intercept | -6.59 | 0.18 | -36.353 | **<0.0001** |
| SW | 0.20 | 0.06 | 3.192 | **0.0018** |
| Sex | -0.67 | 0.13 | -5.450 | **<0.0001** |
| *First time spawners* | | | | |
| No. mates ~ Sex + SW | | | | |
| Intercept | -6.54 | 0.19 | -34.985 | **<0.0001** |
| SW | 0.20 | 0.07 | 3.021 | **0.0030** |
| Sex | -0.69 | 0.13 | -5.350 | **<0.0001** |
